# NiftyTorch: A Deep Learning framework for NeuroImaging

**DOI:** 10.1101/2021.02.26.433116

**Authors:** Adithya Subramanian, Haoyu Lan, Sankareswari Govindarajan, Lavanya Viswanathan, Jeiran Choupan, Farshid Sepehrband

## Abstract

We present NiftyTorch a Deep Learning Framework for NeuroImaging. The motivation behind the development of such a library is that there are scant amount of centralized tool for deploying 3D deep learning for NeuroImaging. In addition, most of the existing tools require expert technical knowledge in Deep Learning or programming, creating a barrier for entry. The goal is to provide a one stop package using which the users can perform classification tasks, Segmentation tasks and Image Transformation tasks. The intended audience are the members of NeuroImaging who would like to explore deep learning but have no background in coding. In this article we explore the capabilities of the framework, the performance of the framework and the future work for the framework.

## I. Introduction

NeuroImaging is a sub-field of Neuroscience where the internal structure of the area interest in an organism is captured. The Images are captured using techniques such as Magnetic Resonance Imaging, Positron Emission Tomography and Computed Tomography. The Image captured in the process has helped the Physicians to diagnose fatal disease such as cancer, Alzheimer’s disease, stroke and several other neurological disorders. The diagnosis although now being a possibility demands expertise from the Physician and is often difficult even for well-trained Physicians to identify these diseases. It is seen that use of deep learning has seem to alleviate these problems for several applications. Deep learning could assist Physician in diagnosis, feature extraction, disease monitoring, reporting, future outcome prediction and so on. Hence, we are building NiftyTorch to accelerate the usage of deep learning in this field. NiftyTorch is pip installable (pip install niftytorch) and the online documentation can be accessed via https://niftytorch.github.io/doc/.

### A. Why NiftyTorch?

There has been an explosive surge of improvements in the field of deep learning in areas such Computer Vision, Natural Language Processing and Speech Recognition whereas a scant amount of developments can be seen in the application of deep learning in NeuroImaging which can be easily transferred from the aforementioned fields.

We identified that one of the main reasons behind such a backlog in the developments across fields is the lack of programming and deep learning knowledge that acts as barrier in imbibing the recent advancements in one area of deep learning to another one. Hence, we want have built NiftyTorch such that it would less than 10 lines of code to train a deep learning on your favourite dataset.

### B. What can NiftyTorch do?

NiftyTorch is built on top the python library Pytorch [1] which gives us flexibility to code complex research models available at ease. NiftyTorch builds on top the code from Pytorch to develop a custom dataloader that can work .nii and .nii.gz format while maintaining the performance.

The library also has state of the Neural Networks for 3D data such as AlexNet, VGGNET, ResNet, ShuffleNet, SqueezeNet and Xnor-Net. Nevertheless the user can develop their own network using the 3D Convolution Building Blocks such as Binary Activation, Channel Shuffle, Fire Module. We also provide Segmentation Networks such U-Net, V-Net and HyperDenseNet.

Our main contribution towards the accelaration of use of deep learning the NeuroImaging field would be the use of Automatic Hyperparameter Tuning. It is our hunch that many research ideas get dropped during empricial analysis due to lack of effective ways to improve performance. We feel that automatic hyperparameter tuning will help neuroscientists cross this barrier and present novel research using deep learning.

We go in detail about what NiftyTorch has to offer in the Framework section.

## II. Related Work

While we offer a solution to the problem of applying deep networks to the field of NeuroImaging there are other solution which also offers different features to solve the same problem. In this section we go about the different such solutions.

Monai framework is a PyTorch-based framework for deep learning in healthcare imaging. It provides workflows for using domain optimized networks, loss functions, metrics and optimizers. Similar to NiftyTorch Monai also supports image loader for loading nifty files and distributed training. It is also optimized for CUDA support.

MedicalTorch is a python based library was one the early libraries in this area it provides datasets, data loader, metrics, models and loss functions.

## III. Features

In this section we describe the features of the package. Fig. 1 is the diagrammatic representation of NiftyTorch architecture. The complete architecture includes 6 modules: NiftyTorchPrep, DataLoader, Transformation, Model, Trainer and Predicotr. We will describe each module’s details in the following text.

**Fig. 1:**
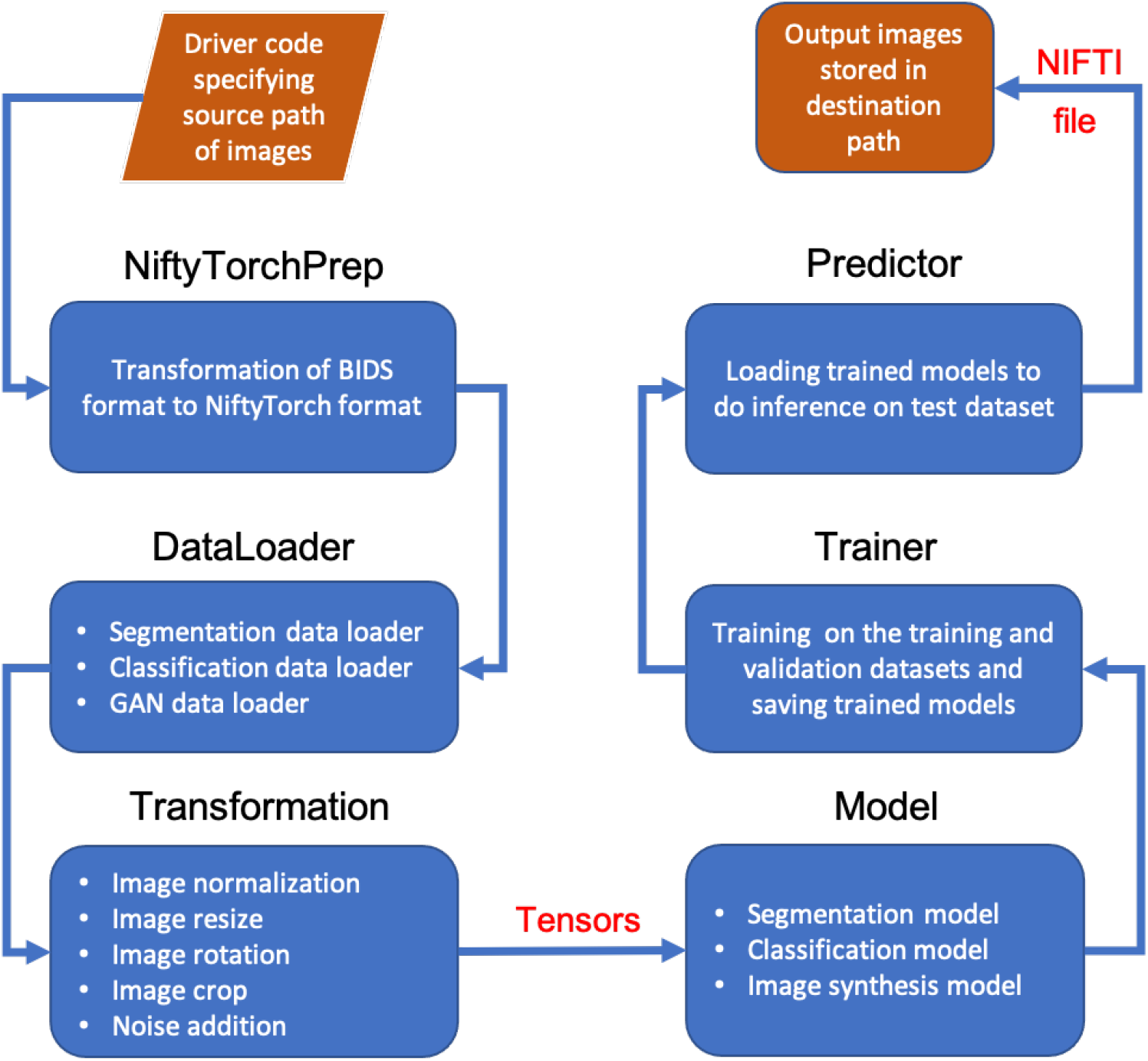
NiftyTorch architecture.

### A. NiftyTorchPrep

NiftyTorchPrep is the dataset structure transformation command to check if BIDS (Brain Imaging Data Structure) [2] format of user’s dataset is correct and transform it into the format that is coherent with data structure used in NiftyTorch. Fig. 2 shows the dataset format transformation after using below niftytorchprep command:

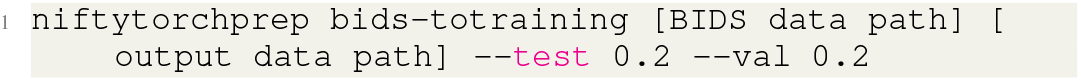

**Fig. 2:**
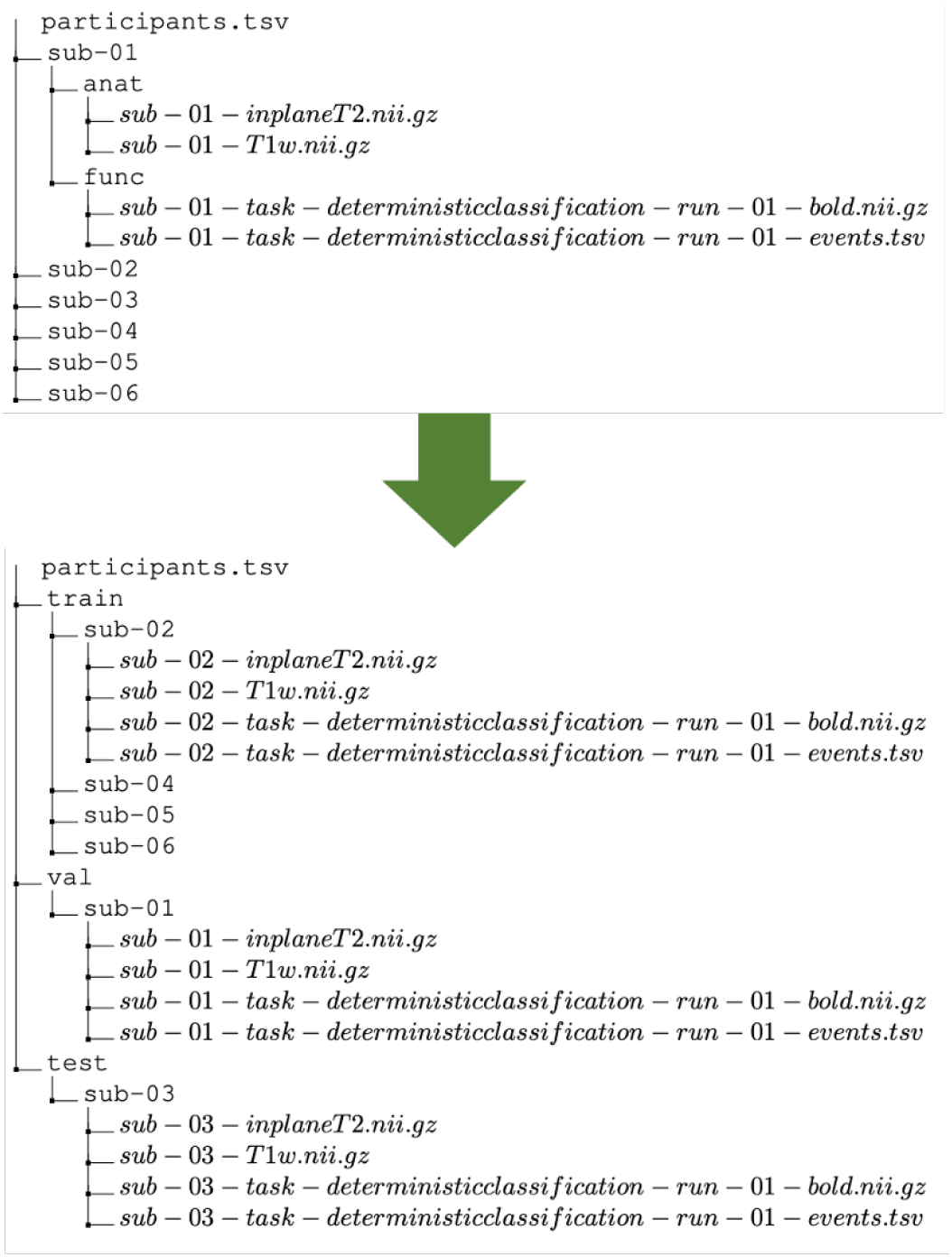
Format transformation from BIDS format to NiftyTorch format.

### B. DataLoader

Data loader is the data loading module of the library and supports three different tasks: classification, segmentation and image transformation tasks. The data loader is built on top of the Pytorch as a result it supports multiple workers, batch data loading and applying transformations.

We have changed the internal working of the loader to support nifty files i.e. it essentially supports all the files supported by nipy library. The data loader in NiftyTorch supports pre-processing such as resize, rotation and identifying the area of interest.

The NiftyTorch has three types of Data Loaders:

- Classification DataLoader
- Segmentation DataLoader
- Paired DataLoader
- Image transformation DataLoader

#### 1) Classification Data Loader

Classification Data Loader as seen in the name is used for classification tasks. The built in classification networks made available in the NiftyTorch have also been integrated with the classification data loader. The Classification Data Loader requires the data to be formatted in the following:

1. Train
  a. subject name 1
    - t1w.nii.gz
    - t2w.nii.gz
  b. subject name 2
    - t1w.nii.gz
    - flair.nii.gz

Similar directory distribution is required for the validation dataset.

The labels for each of the dataset is provided in the form of a csv. The csv file must contain two columns one for the subject name and the other one is for label.

An example for a the csv file would be as shown in the Table 1.

Along with the label, the users can also provide demographic data which is categorical in nature as seen in Table 1. These features are included inside the neural network just before we use reshape the input tensor to 2D.

The classification data loaders allows to load multiple modalities of the same patient together so that we can leverage as much data as possible. The modalities are stack on the channels on top of the other.

| Subject 1 | Label | Age |
| --- | --- | --- |
| subject name 1 | benign | 22 |
| subject name 2 | malign | 77 |

A sample snippet of defining a Classification Data loader can be seen below.

**Listing 1:**
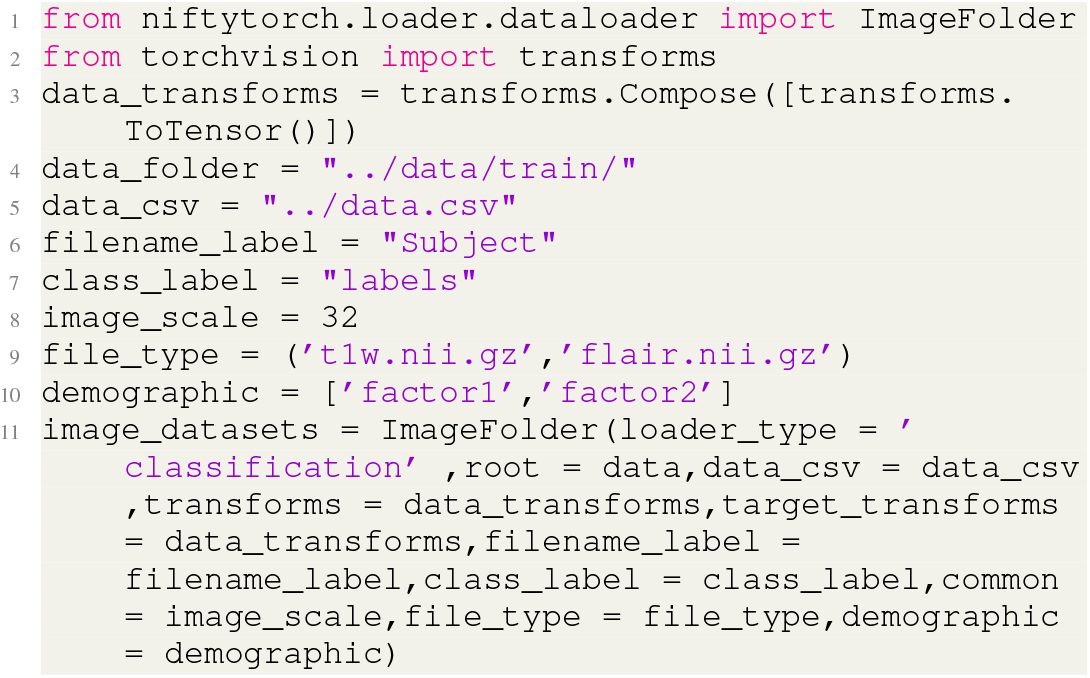
Classification loader example

#### 2) Segmentation Data Loader

Segmentation data loader is used for segmentation tasks. The Segmentation data loader has been integrated with the segmentation networks U-net, V-net and HyperDenseNet. The directory distribution of files are done as follows:

1. Train
  a. subject name 1
    - t1w.nii.gz
    - seg.nii.gz
  b. subject name 2
    - t1w.nii.gz
    - seg.nii.gz

The segmentation loader supports region of interest extraction which is very crucial for segmentation tasks since the output masks are generally sparse in nature. This region of interest pre-processsing is not needed for validation set as we want to infer on the entire image. Hence, it can be removed from the workflows with ease.

Another important factor is the multi-dimensional mask for prediction as generally there might be different types of labels for the same tumor in the brain. This is also supported by the NiftyTorch.

A sample snippet of defining a segmentation Data Loader can be seen in Fig 2.

**Listing 2:**
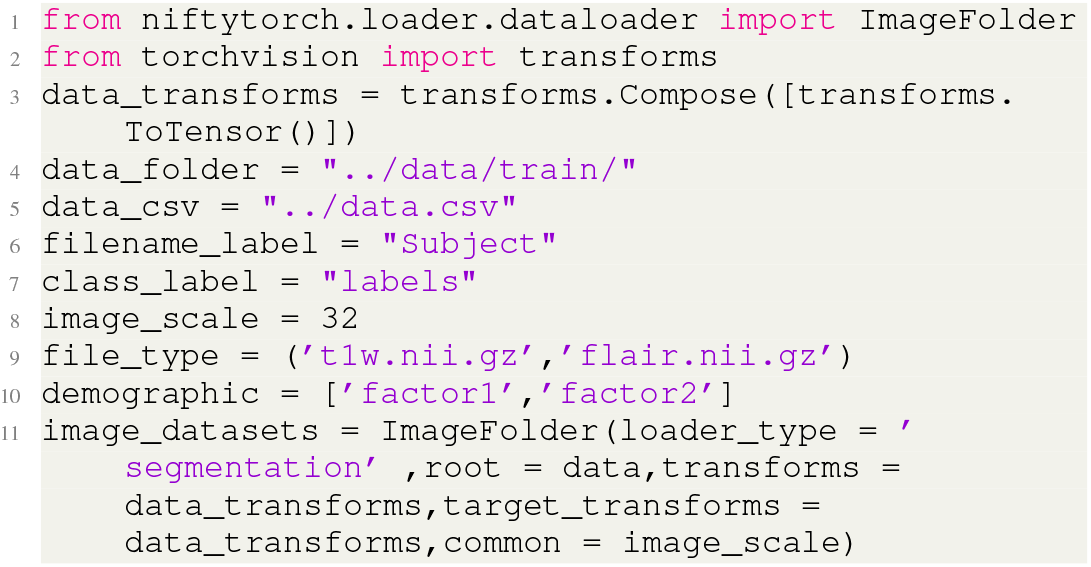
Segmentation Loader example

#### 3) Paired Data Loader

In NeuroImaging we always see that the Data is a few i.e. of orders 1000 samples. The usual networks like ResNet and AlexNet will have a hard time learning from these samples.

Hence, we need metric learning loss functions which will improve the performance of the deep learning models. Usually the metric learning loss functions require multiple data points to act as anchors and negative samples. This is currently supported by NiftyTorch we believe that this will help neuro-scientists to work on critical applications with scant amount.

The one of the functionalities supported by the Data Loader is the number of negative samples to be sampled by the Data Loader which is needed for implementation such as Angular Loss, Circle Loss etc,.

The Directory structure of Paired Data Loader is same as that of the Classification Data Loader.

A sample snippet of defining a Paired Data Loader can be seen in Fig 3.

**Fig. 3:**
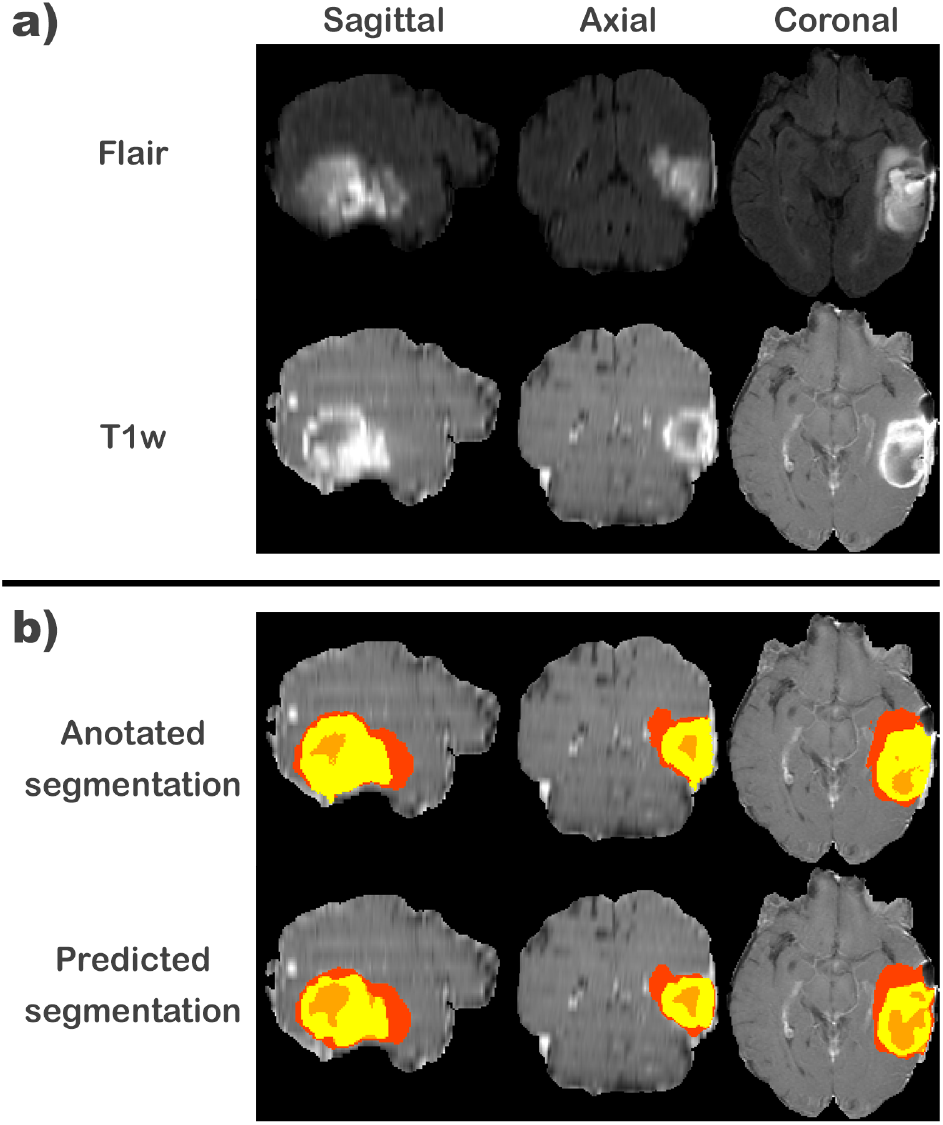
multi-modal multi-class Tumor segmentation example, using BRATS dataset.

**Listing 3:**
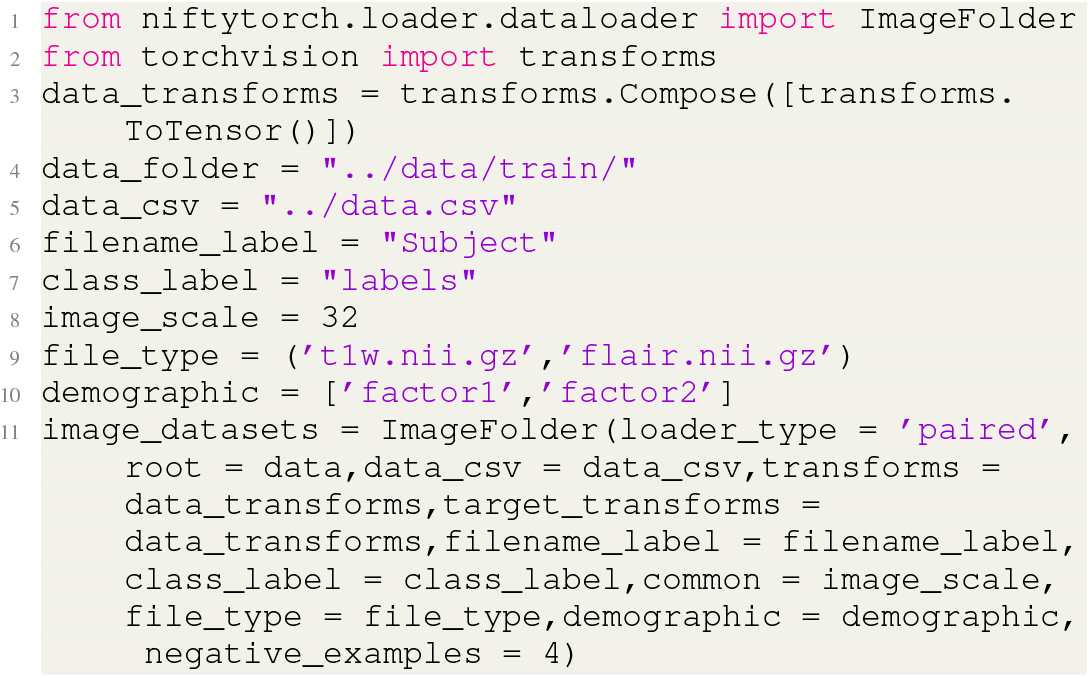
Paired loader example

#### 4) Image transformation Data Loader

Image transformation is a useful technique validated by deep learning methods for recent years. It includes medical image reconstruction from k-space, image super-resolution, noise reduction and image synthesis. NiftyTorch supports image transformation tasks by GAN (Generative Adversarial Network) [3] and specified GAN training dataloader. the directory structure of image transformation DataLoader is the same as that of the classification DataLoader.

**Listing 4:**
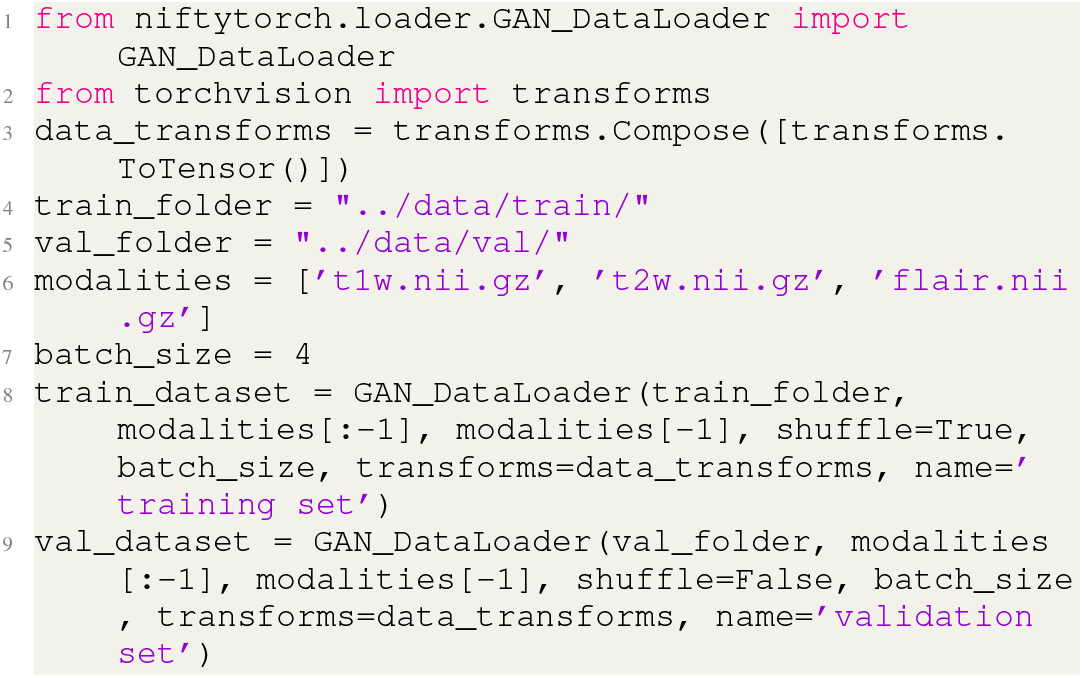
GAN DataLoader example

### C. Transformation

Transformation module is used to do the image processing and data augmentation before the training stage. Data Augmentation is made available for the users to increase the data as it is usually less in compared to traditional deep learning problems. The different types of preprocessing and data augmentations available in NiftyTorch are as follows.

#### 1) Normalization

Min-max normalization and Z score normalization are supported by NiftyTorch.

#### 2) Rotate

This operation allows the user to rotate the tensor 90 degree, 180 degree and 270 degrees along different axis.

#### 3) Noise Addition

This operation allows the user to add to input tensor which can help us to improve next performance in low data scenario and as well as when the data is sparse.

**Listing 5:**
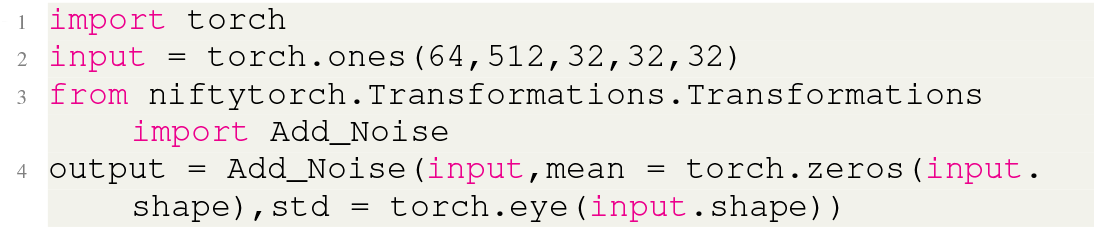
Noise Addition example

#### 4) Random Segmentation Crop

This operation allows the user to randomly crop areas of the input image around ground truth segmentation masks.

**Listing 6:**
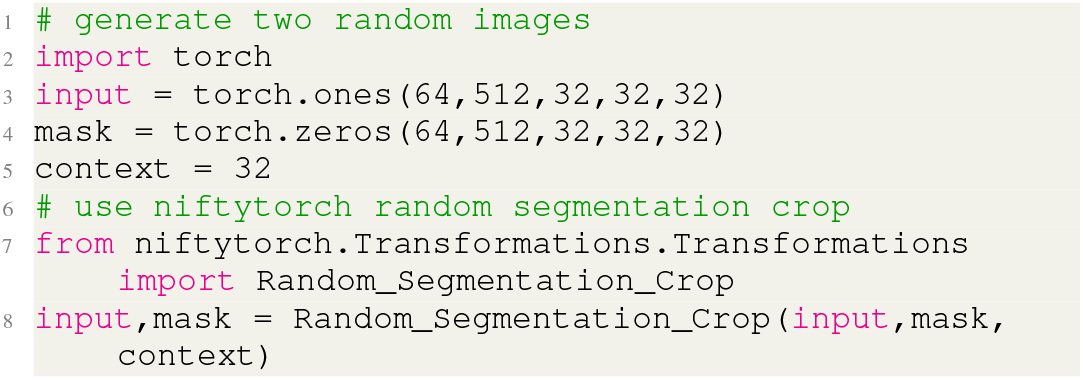
Random Segmentation Crop example

#### 5) Resize

The Resize operation resizes the input tensor and mask tensor if it present to the provided size using interpolation.

**Listing 7:**
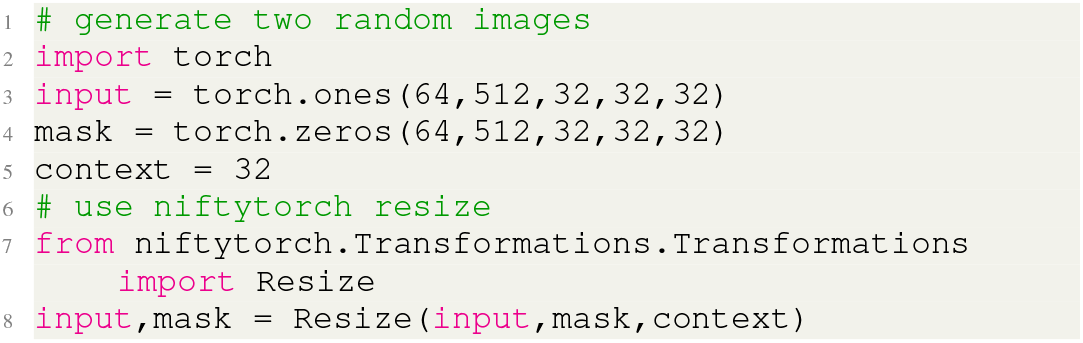
Resize example

### D. Models

The NiftyTorch has a collections of models which can be used directly with less than 3 lines of code. In the section below we brief about the changes we have made to adopt the models for 3D data, the tunable hyper-parameters of the model and the ones which are considered for the automatic hyper-parameter tuning. All the models have support for demographic data.

#### 1) AlexNet

In NiftyTorch for AlexNet [4] we only fix the architecture of the model and rest of the hyperparameters are tunable. The output dimension of the matrix dimension is the tunable by the user and the input size of the network is not fixed.

**Listing 8:**
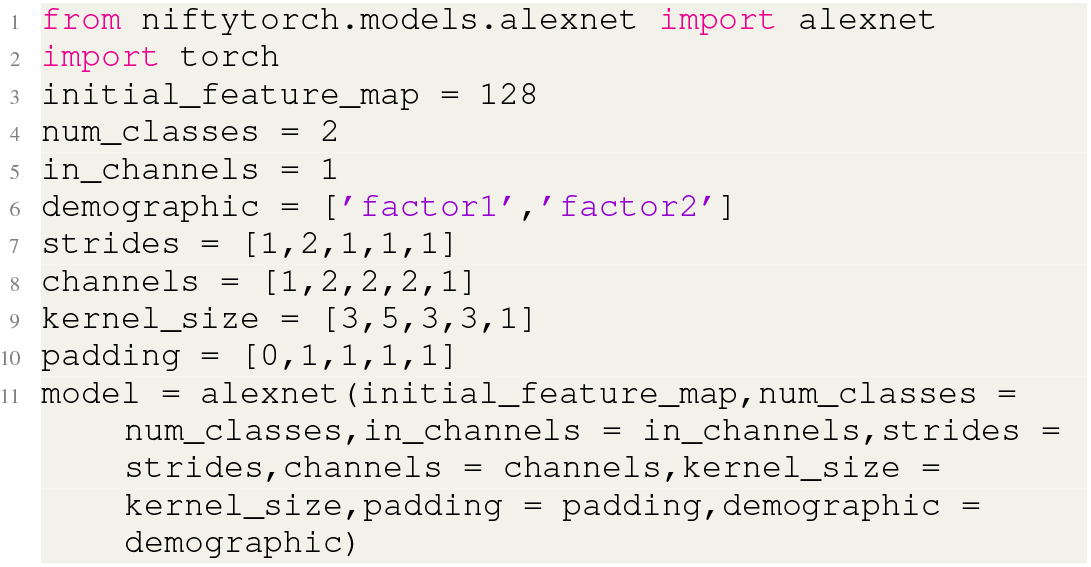
Alexnet example

#### 2) Vgg Net

In NiftyTorch, Vgg Net [5] we have added support for the different versions of VGG Network, this include A,B,D and E. The VGG also has Batch-Normalization included in it to boost the performance of the network.

The users can also change the structure of the network to have a custom version.

**Listing 9:**
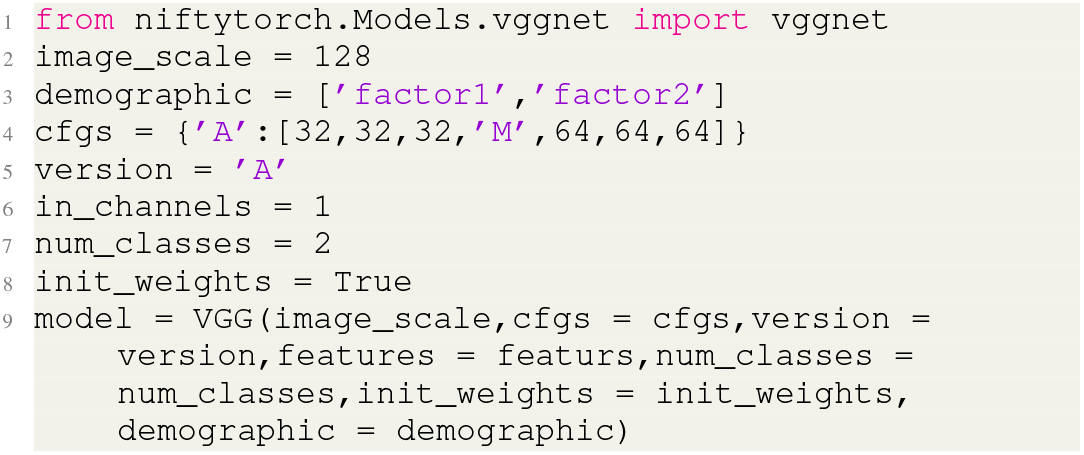
VGG Net example

#### 3) ResNet

ResNet [6] being one of the most powerful networks we also provide support for it. In Resnet we include two type of internal structure one is the bottleneck and basic block. Unlike the other networks in NiftyTorch Resnet also allows tuning of the number of layers in the network. Although a hyper-parameter this is not included in automatic hyper-parameter tuning as this causes a lot of parameter in the framework.

**Listing 10:**
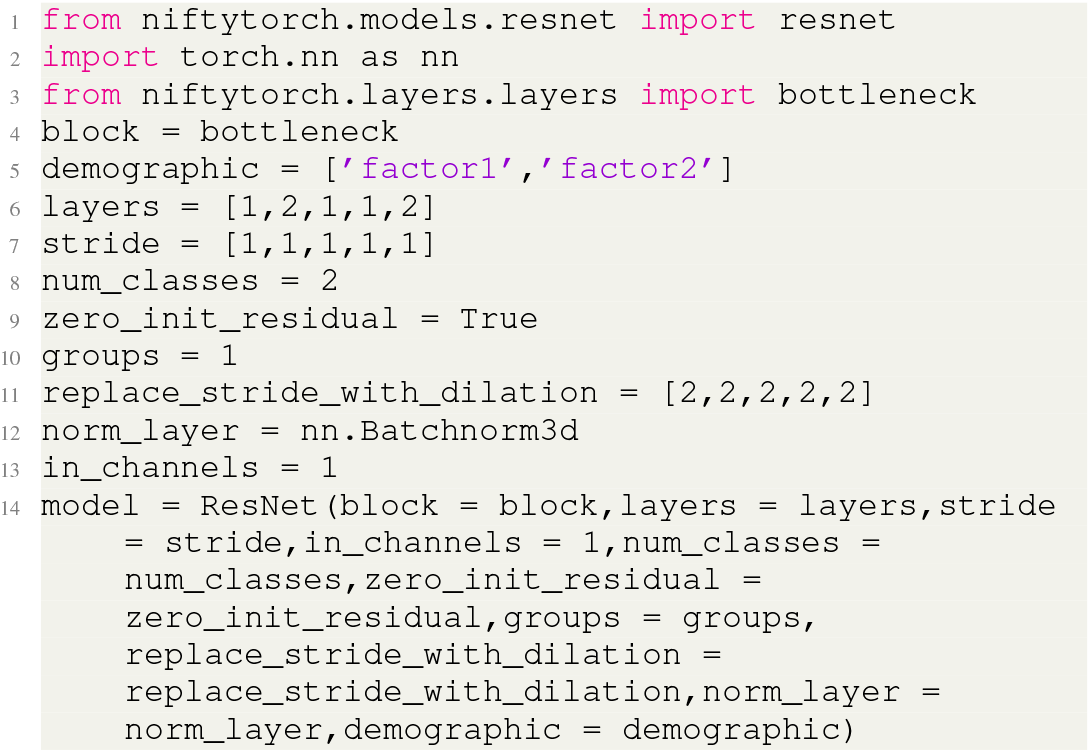
ResNet example

#### 4) SqueezeNet

Considering that the data size is large and occupies a lot of memory we believe that it is important to have support networks which do not occupy as much memory as other models. Hence, we need squeezenet [7] which helps us attain high accuracy as g as Alexnet with fewer parameters.

In NiftyTorch we support SqueezeNet version 1.0 and 2.0. In terms of Automatic Hyper-parameter we currently do not support Hyper-parameter tuning.

**Listing 11:**
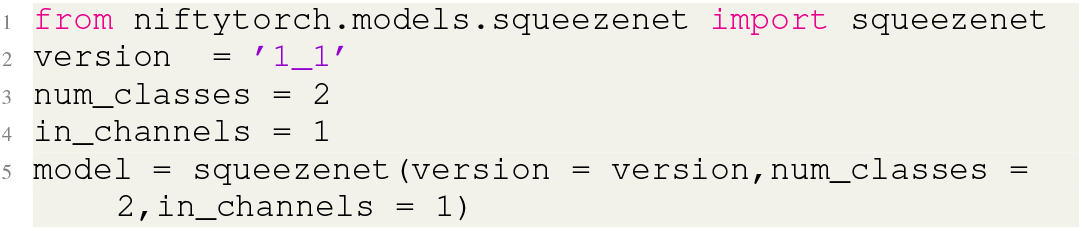
SqueezeNet example

#### 5) ShuffleNet

Again considering the memory efficiency issues in the Neuroscience, NiftyTorch supports ShuffleNet [8].

**Listing 12:**
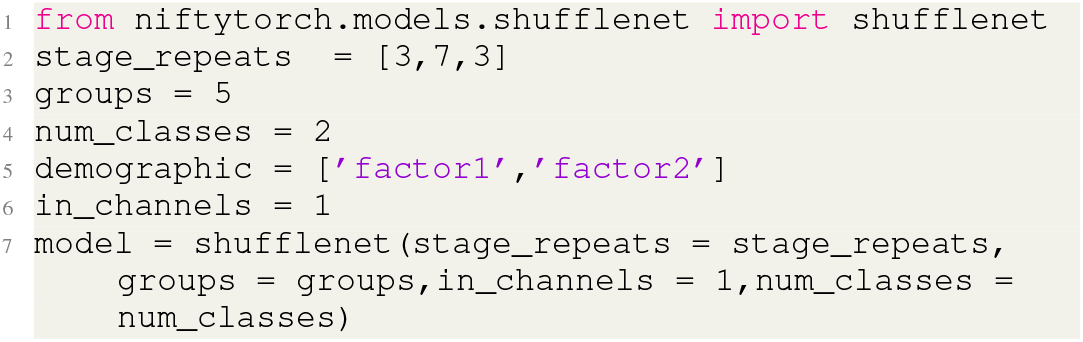
Shufflenet example

#### 6) Xnor Net

Considering the huge tensor size, the memory and speed both become an issue we have implemented Xnor Net [9] in NiftyTorch to counter these aspects.

In NiftyTorch we’ve implement XNOR Net with an Alexnet base.

**Listing 13:**
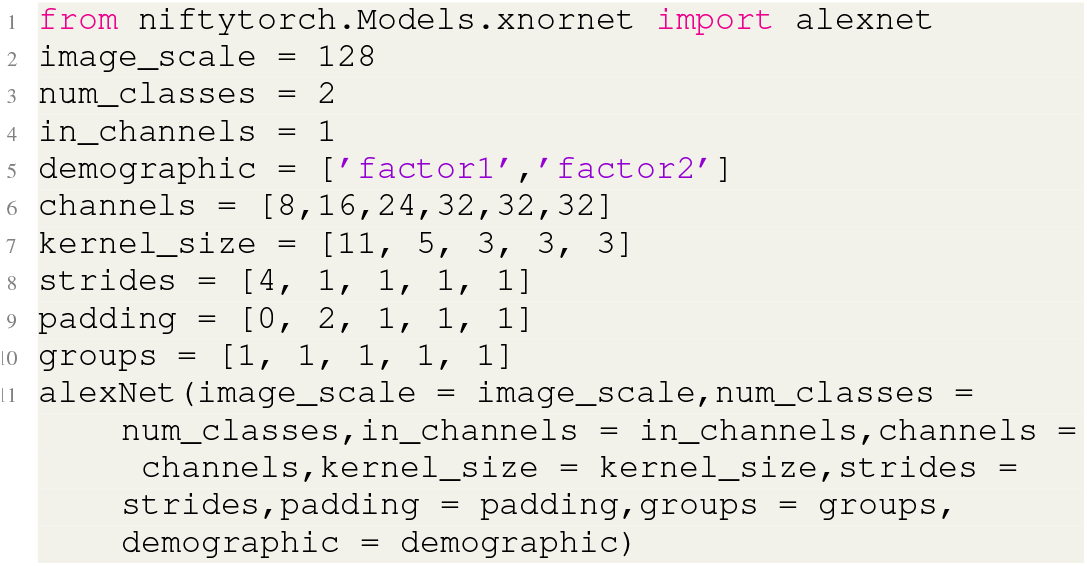
Xnor net Example

#### 7) U-Net

The image segmentation models UNet [10], VNet [11] and HyperDenseNet [12] allow the users to train model, store and make use of an already trained model, segment unseen images. The model supports multiple modality inputs and can generate both single and multiple channel outputs. Users can tune the model as per their requirement. In terms of compute, parameters like ‘cuda’, ‘num workers’, ‘device ids’ can help train the model with minimal consumption of resources. In terms of results accuracy(model performance), ‘stride’,’padding’,’kernel size’,’downsample’ parameters can be made use of.

Based on the test results, we derived inferences on the behavior of the three models. Apart from the close differences in the architecture, U-Net seems to be perform well in predicting more accurately when compared to V-Net. Whereas, HyperDenseNet is best suitable for image samples of very low dimensionality.

Unlike the other models U-Net is built for the purpose of segmentation, this U-Net supports multi-class segmentation. A resulting segmentation and comparison can be seen below.

For an image of voxel dimension 138*170*137 the parameters are as seen in the snippet below:

**Listing 14:**
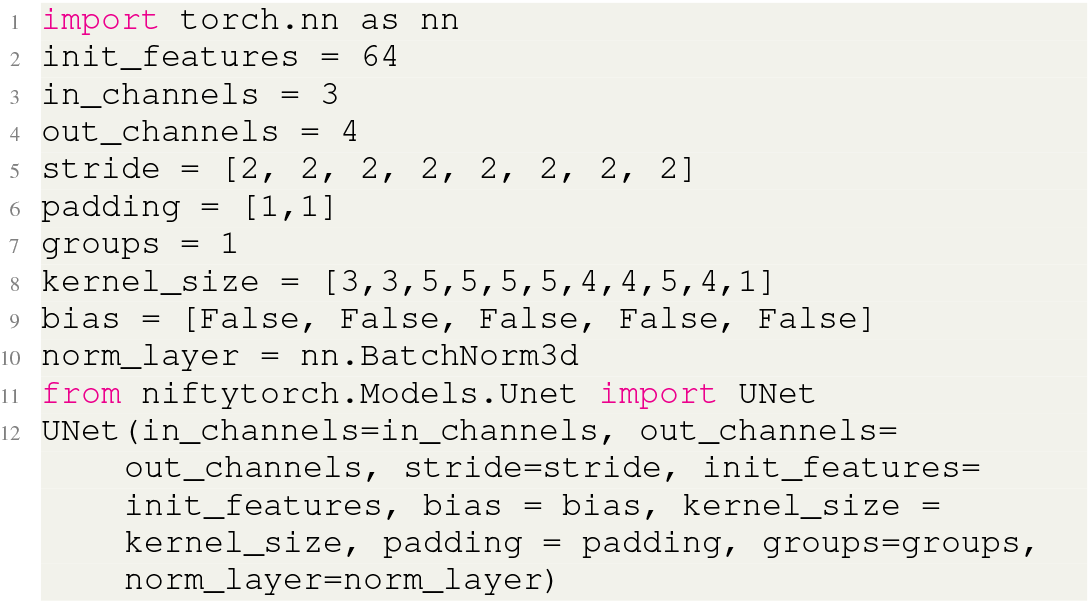
U net Example

#### 8) V-Net

Similar to U-Net NiftyTorch also supports to V-Net.

#### 9) HyperDense-Net

Similar to U-Net NiftyTorch also supports to HyperDense-Net. HyperDense-Net was based on DenseNets and extended a better performance for multi-modal segmentation.

#### 10) Pix2pix

NiftyTorch also supports neuroimage synthesis tasks. Pix2Pix [13] is a well developed image transformation algorithm and we extended 2D Pix2Pix model to 3D to handle neuroimage modalities synthesis.

#### 11) SC-GAN

In order to fully support neuroimage synthesis tasks, NiftyTorch incorporates multi-modality neuroimaging synthesis algorithm SC-GAN [14].

Pix2pix and SC-GAN share the same defination code, here we only show one snippet about how to define SC-GAN.

**Listing 15:**
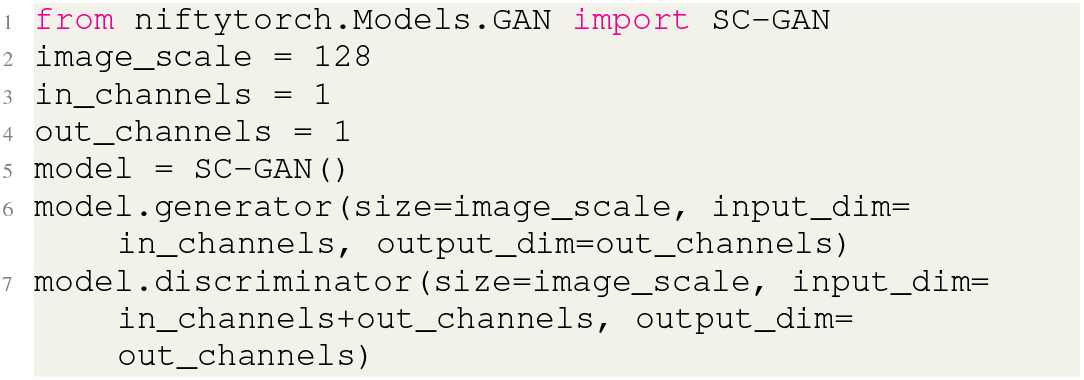
GAN Example

### E. Trainer and predictor

NiftyTorch supports modularized training and predicting procedure. In order to simplify the training and predicting processes for the user friendly purpose, NiftyTorch enables two modules called trainer and predictor. Users only need to specify generic training parameters like batch size, training epochs, learning rate etc and pass the predefined models to trainer. The trainer will finish the training process on training and validation dataset automatically and save the trained model in the user defined model-saving path. Predictor fetches trained model and do the inference on test dataset and save output images in the user defined image-saving path.

## IV. Model customization

We are not constrained with the models presented in the above section. We’re able to extend building networks beyond the conventional network as we make the inner building blocks available which we will be described in section 3.3.

We have also made a demo available on how to use a custom network with DataLoader mentioned in Section 3.1. It also supports adding attention mechanism and demographic data.

### A. Convolutional Building Blocks

The Convolutional Building Blocks are the smallest unit used in the neural network which act as backbone of the building block. We can mix and match each of the Convolutional building block to form our custom models.

#### 1) BottleNeck Unit

BottleNeck is the building block of the residual neural network which is used in highly deep network.

**Listing 16:**
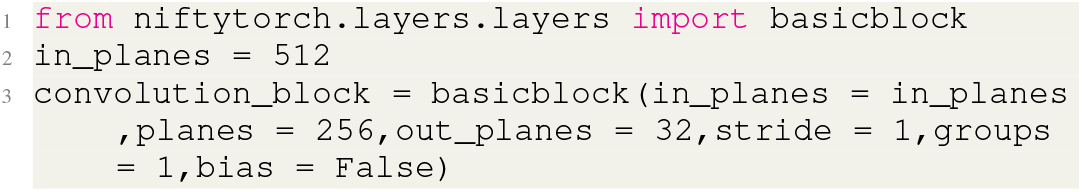
BottleNeck unit example

#### 2) Shuffle Unit

ShuffleNet unit specially designed for small networks with similar basis to residual building block. Channel shuffle operation makes it possible to build more powerful structures with multiple group convolutional layers. The convolution is a combination of depth-wise convolution and pointwise convolution.

**Listing 17:**
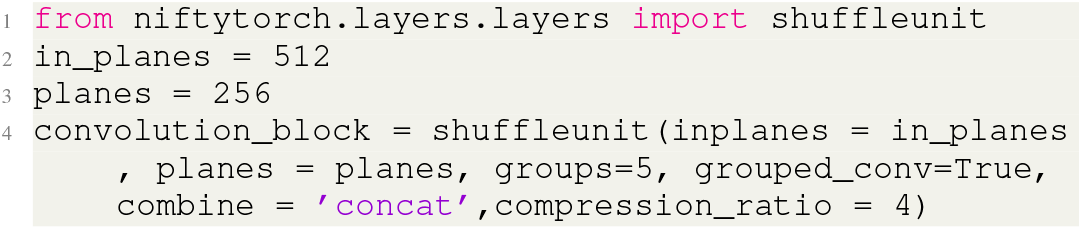
Shuffle unit example

#### 3) Fire Module

A Fire module is comprised of: a squeeze convolution layer (which has only 1×1 filters), feeding into an expand layer that has a mix of 1×1 and 3×3 convolution filters. It is the foundation of SqueezeNet.

**Listing 18:**
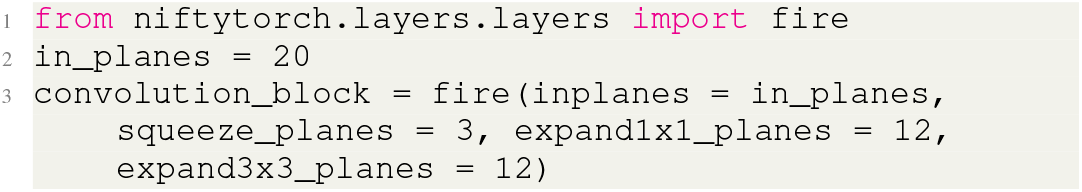
Fire Module example

#### 4) Binary Activation

The Binary Activation is a module which speeds up the computation of activation through binarization the feature map.

**Listing 19:**
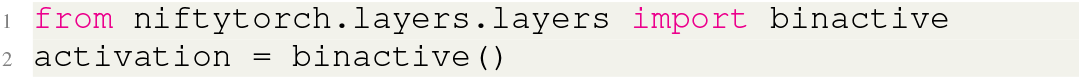
Binary Activation example

#### 5) Binary Convolution

The Binary Convolution is Binarized formulation of the convolution operator which as a results reduces the computation time and the storage space.

**Listing 20:**
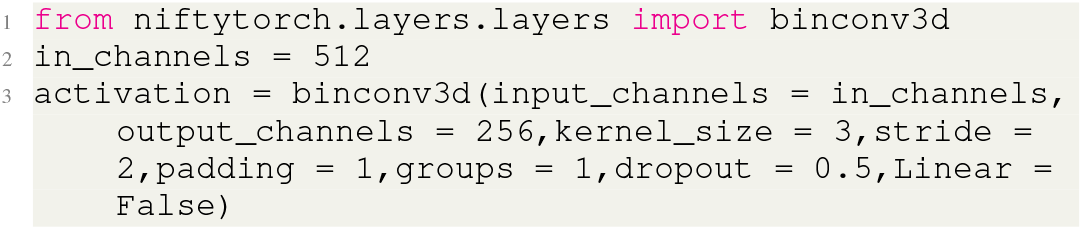
Binary Convolution example

### B. Loss Functions

The Loss Functions are made such that it is easily integratable with custom models as well as the current models.

#### 1) Cross Entropy

In NiftyTorch models the default loss function is CrossEntropy with equal weights. The different weights are supported but it needs to be passed explicity. The snippet below shows below on how to use explicity use weighted cross entropy.

**Listing 21:**
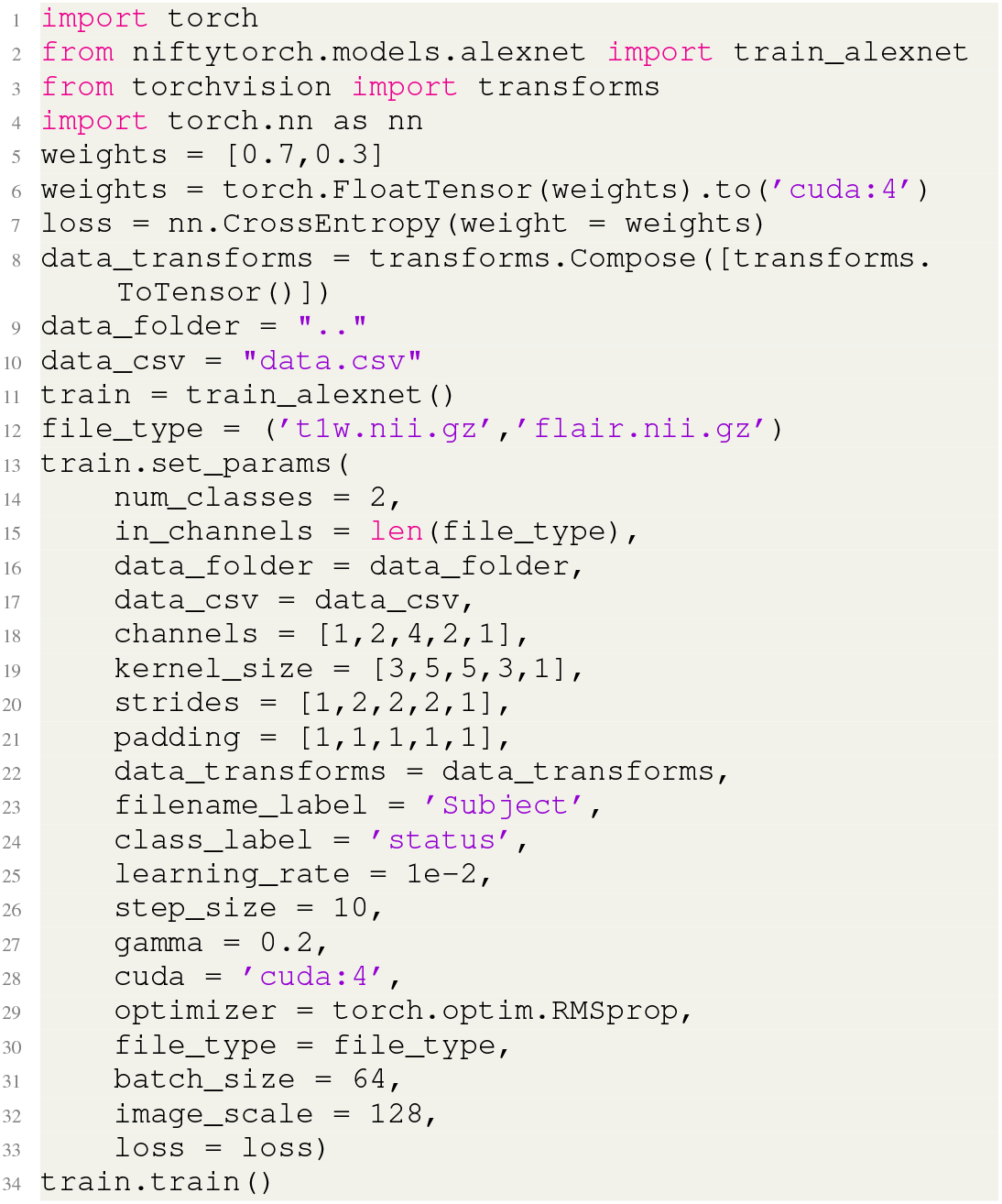
Cross Entropy Loss example

#### 2) Focal Loss

The default parameters to the Focal Loss [15] are 0.5 weights for each and exponent is 2. Unlike Pytorch’s implementation of the CrossEntropy Loss both the weights and exponent can be simple list and float respectively

**Listing 22:**
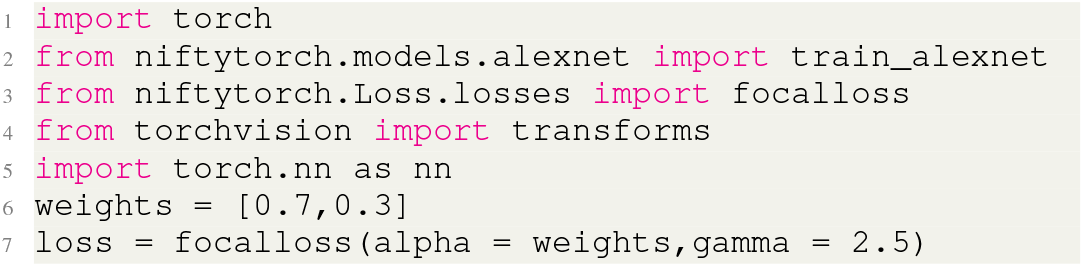
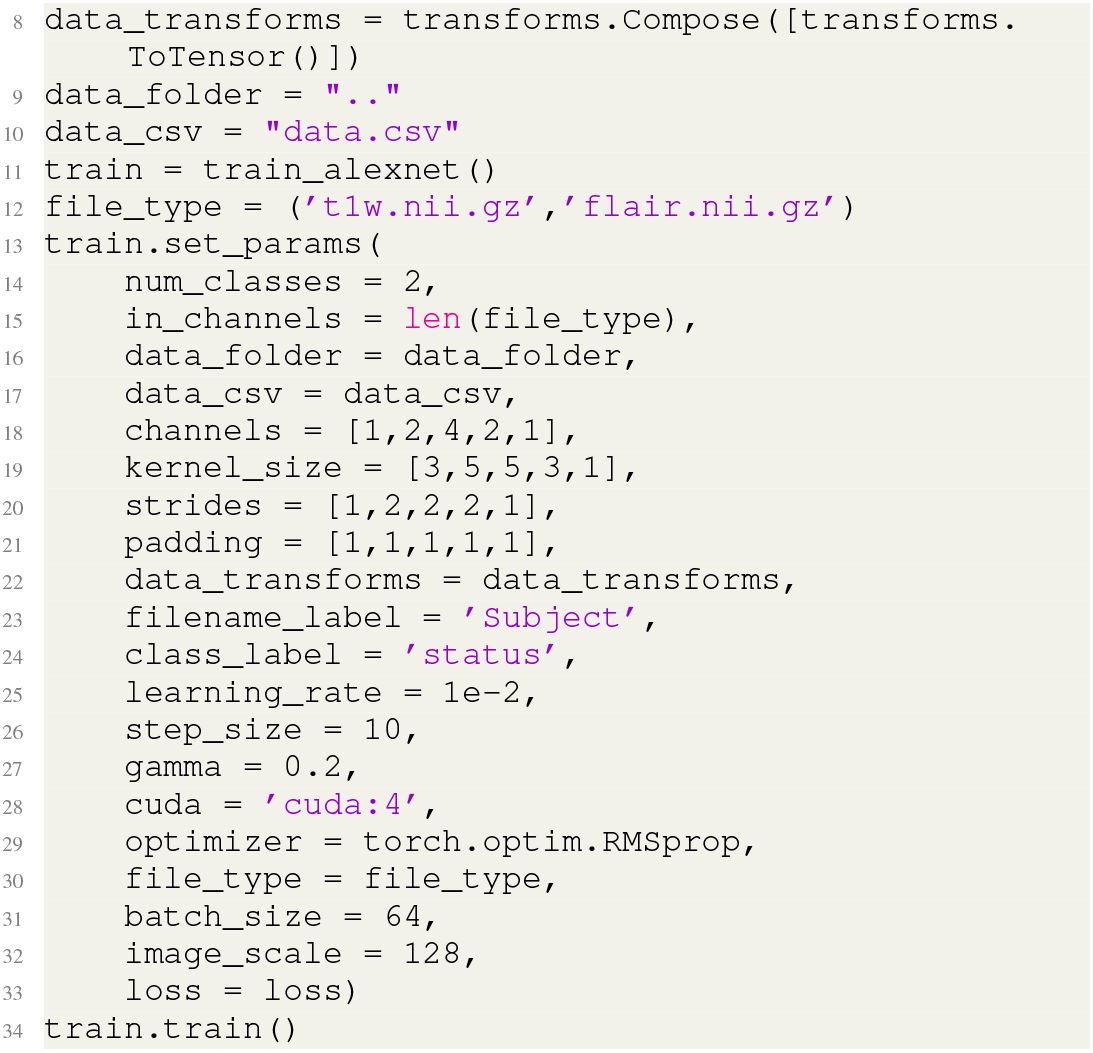
Focal Loss example

#### 3) Focal Dice Loss

Focal Dice Loss can be used imitate Dice Loss by setting beta = 1.

**Listing 23:**
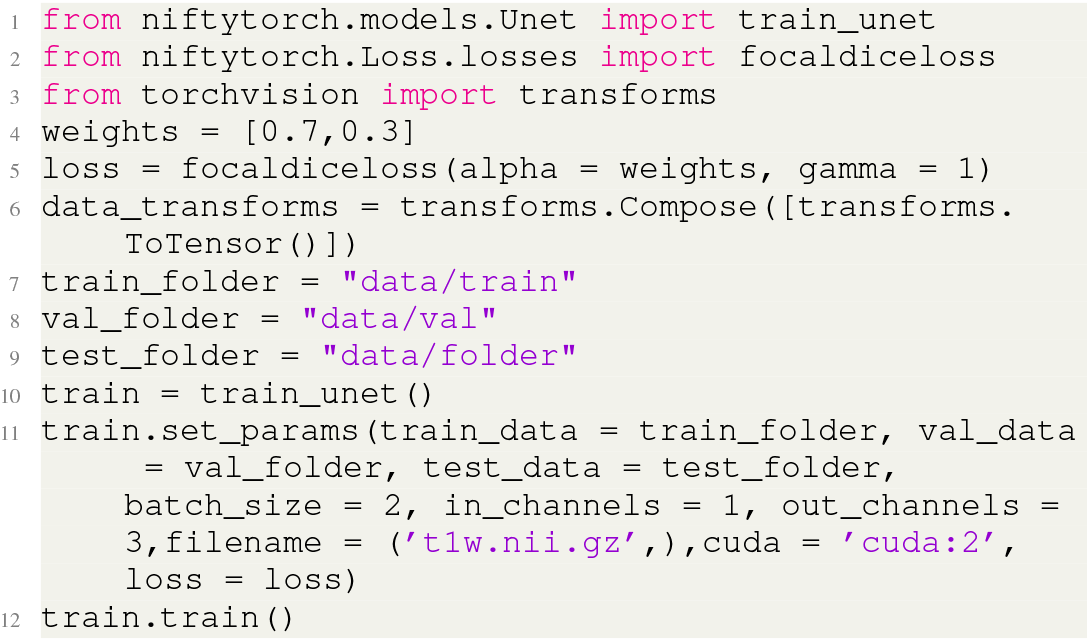
Focal Dice Loss example

#### 4) Lovasz Softmax

Lovasz softmax [16] loss is used for handling unbalance-class segmentation problem.

**Listing 24:**
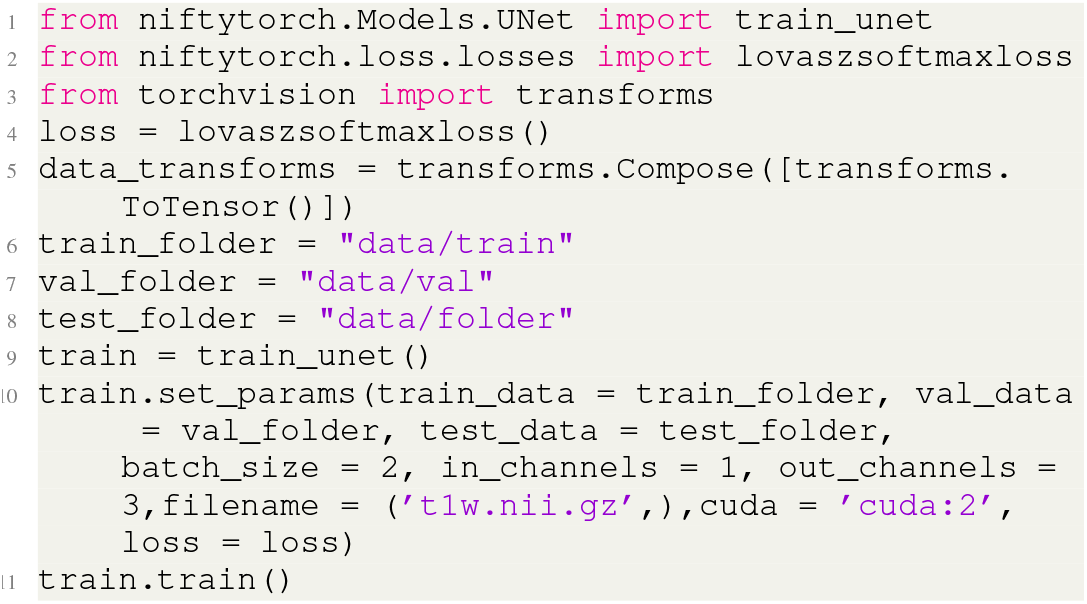
Lovasz Softmax example

#### 5) Adversarial loss

Adversarial loss is used for GAN (generative adversarial nets) [3] training process. NiftyTorch supports GAN related algorithm Pix2Pix and SC-GAN.

#### 6) Soft N Cut Loss

Soft N Cut Loss is used for unsupervised image segmentation.

**Listing 25:**
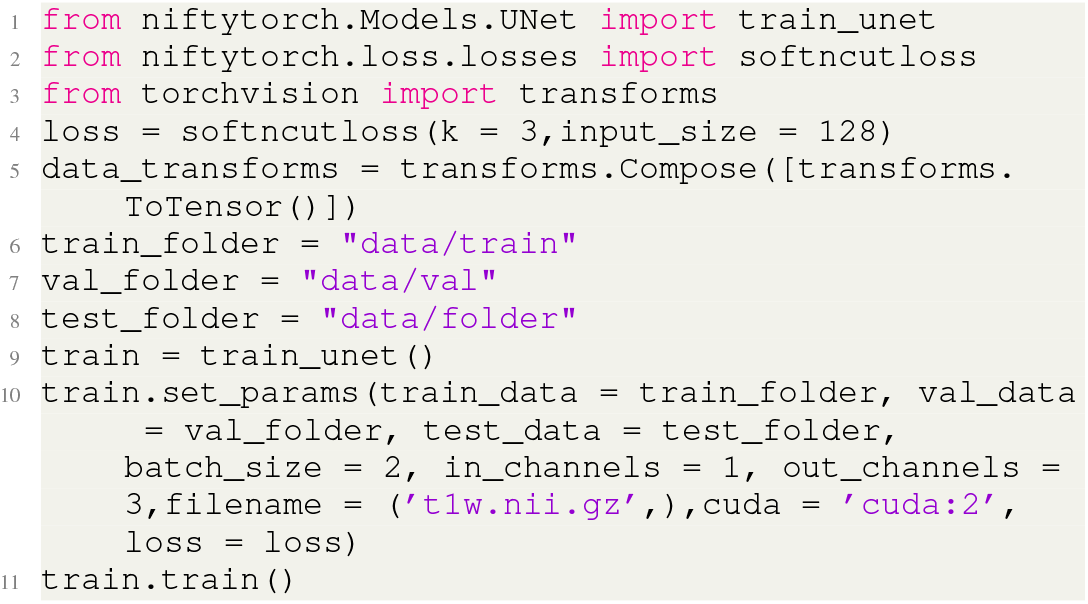
Soft N Cut Loss example

### C. Attention Mechanism

Attention Mechanism is of high interest in the area of NeuroImaging where the demographic data and other forms image data is of equal importance to the MRI data. Hence, we support two types of Attention Mechanism.

#### 1) Positional Attention

Positional Attention [17] attends on the positional features i.e the (x,y) across the channels. It is particularly effective in the cases where the input feature map distribution has low spatial variance.

**Listing 26:**
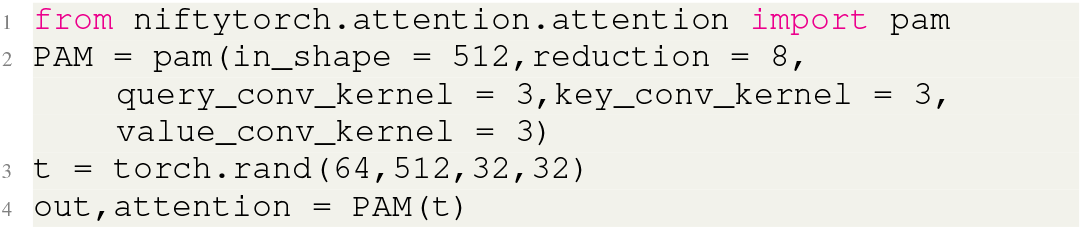
Positional Attention example

#### 2) Channel Attention

Channel Attention [17] attends on the channels features across the (x,y) positions. It is particularly effective in the cases where the input channel distribution has low variance.

**Listing 27:**
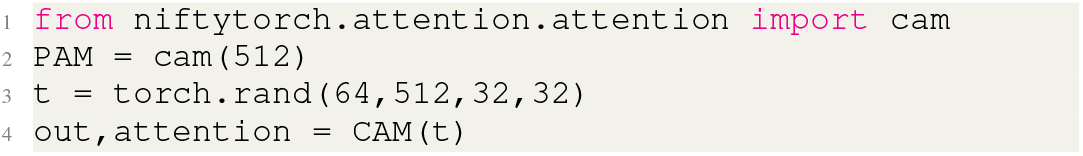
Channel Attention example

### D. Training

In NiftyTorch we support several different types of training mechanism to put the user at ease irrespective of the type of Task.

#### 1) Data level Parallelization

The Data Parallelization mechanism allows the user to shift the data across multiple GPUs and place the model on GPU for training. From a performance stand point we suggest the user to use this when the data is too large and cannot be used with single GPUs.

#### 2) Multi Scale Training

In several segmentation tasks and some classification task the scale of the input matters especially when input data is sparse in such cases we suggest using Multi-Scale Training made available in NiftyTorch. The Multi-scale Training changes the input image scale using interpolation across each batch to make the model robust to the sparsity. It can be enabled and disabled for validation as per users interest.

#### 3) Automatic Hyperparameter Tuning

One other important features of NiftyTorch is that it can do automatic hyper-parameter which can help researcher to optimize the models such that it optimizes the validation accuracy. This has been seen to save time of the researcher in writing laborious loops for doing hyperparameter tuning.

## V. Demos

In this section, We present the demo results of multi-class tumor segmentation and T2w modality synthesis tasks using NiftyTorch to show the effectiveness of NityTorch.

### A. Multi-class tumor segmentation

Fusing information from multiple modalities and in order to visualize the results more clearly the library provides multi class image segmentation functionality. The labelled input image under which the test was conducted was imbalanced and hence a weighted loss function was used to train the model. Here we show one demo, for which model was trained on BRATS dataset [18].

Fig. 3 shows multi-class tumor segmentation result on the test data. Fig. 3.a) shows the Flair modality and T1w modality. First row of Fig. 3.b) is the annotated tumor segmentation masks by technician and the second row is the tumor segmentation masks generated by trained segmentation algorithm built-in NiftyTorch.

### B. Modality synthesis

For the neuroimage modality synthesis task, we chose T2w modality synthesis using T1w as input modality, which is also the modality transformation from T1w to T2w. Data used for this demo were HCP dataset [19]. NiftyTorch support generative adversarial nets like Pix2Pix and multi-modality neuroimage synthesis algorithm SC-GAN.

Fig. 4 shows the inference results on the test dataset using trained GAN model. The first row of Fig. 4.a) is the input modality T1w, the second row is the target modality T2w and the third row is the synthetic T2w modality generated by trained GAN model built-in NiftyTorch. Fig. 4.b) is the learning curve on the validation data and Fig. 4.c) is the structural similarity index (SSIM) evaluation on the validation data.

**Fig. 4:**
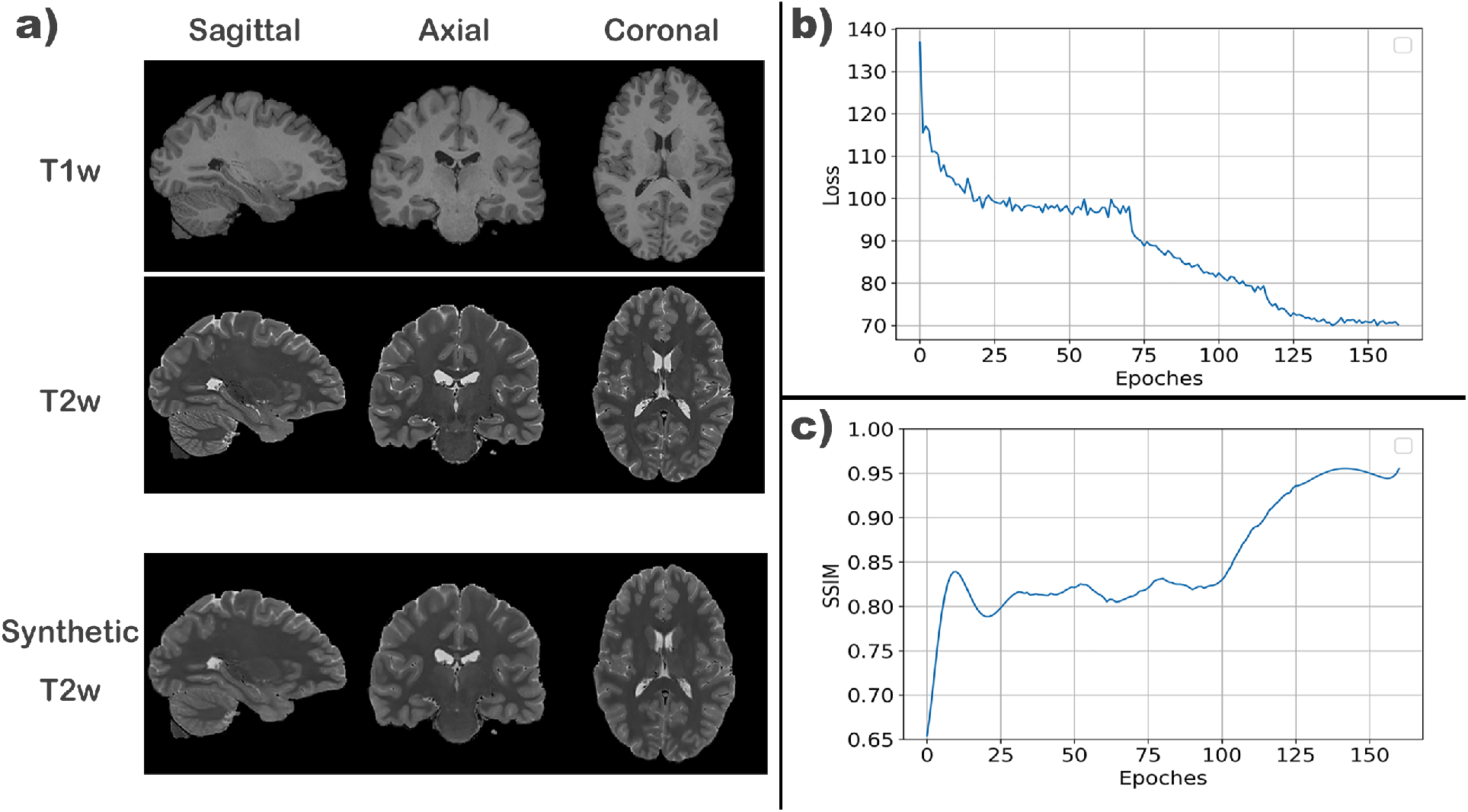
An example output demonstrating T2w synthesis from T1w, using Human Connectome Project 0.7 *mm*^3^ (without downsampling input data).

## VI. Installation Instructions

### A. Prerequisites

NiftyTorch is mainly based on PyTorch, and most of its prerequisites are similar with PyTorch. It has been generally tested over Linux, macOS, and Windows. For Linux users, NiftyTorch requires users to have glibc 2.17 or greater. For macOS users, NiftyTorch requires Yosemite or greater. For Windows users, NiftyTorch requires Windows 7 or greater. To obtain the latest GPU acceleration features, users also need to ensure their CUDA devices are greater than Kepler architecture (i.e. compute capability must be greater than 3.5), and CUDA toolkit is installed properly.

### B. Create a conda environment (recommended for beginners)

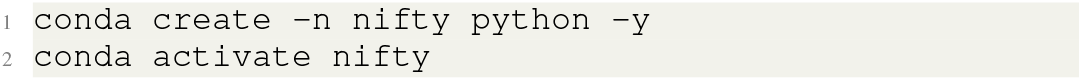

### C. Install NiftyTorch

NiftyTorch can be installed using:

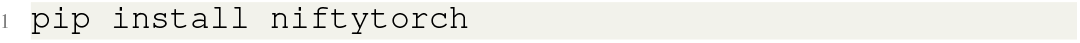

Niftytorch requires the following Python dependencies: torch (greater than 1.4.0) optuna (greater than 1.4.0), torchvision (corresponding to torch), nibabel, numpy (greater than 1.16.4), pandas, nipy, colorlog, alembic, cliff, tqdm, matplotlib, scikit-image. All the requirements will be automatically installed with NiftyTorch.

### D. Visualizing Results

NiftyTorch predict functions would output segmentation or classification results in NIFTI format. Users can utilize any nifti viewer (e.g. FSLeyes) to visualize the output.

## VII. Discussion

We present a deep learning package titled NiftyTorch handling neuroimaging analysis tasks. NiftyTorch has various builtin algorithms and supports customized neural networks by using different builtin building blocks, loss functions, attention modules and user friendly training and predicting pipeline.

## Acknowledgments

This work was supported by the National Institute of Biomedical Imaging and Bioengineering (P41EB015922 and U54 EB020406). We would like to thank Dominik Krzemiński, Sara Morsy and Kaori Lily Ito for developing niftytorchprep plug in (as part of OHBM Hackathon 2020).

## Notes

### Competing Interest Statement

The authors have declared no competing interest.

